# Chemical transcription roadblocking for nascent RNA display

**DOI:** 10.1101/2019.12.26.888743

**Authors:** Eric J. Strobel, John T. Lis, Julius B. Lucks

## Abstract

Site-specific arrest of RNA polymerase is fundamental to several technologies that measure RNA structure and function. Current *in vitro* transcription ‘roadblocking’ approaches inhibit transcription elongation using a protein blockade bound to the DNA template. One limitation of protein-mediated transcription roadblocking is that it requires the inclusion of a protein factor that is extrinsic to the minimal *in vitro* transcription reaction. In this work, we show that interrupting the transcribed DNA strand with an internal desthiobiotin-triethylene glycol modification efficiently and stably halts *Escherichia coli* RNA polymerase transcription. To facilitate diverse applications of chemical transcription roadblocking, we establish a simple and sequence-independent method for the preparation of internally modified double-stranded DNA templates by sequential PCR and translesion synthesis. By encoding an intrinsic stall site within the template DNA, our chemical transcription roadblocking approach enables nascent RNA molecules to be displayed from RNA polymerase in a minimal *in vitro* transcription reaction.

## Introduction

The *in vitro* display of nascent RNA molecules from halted transcription elongation complexes is a powerful tool with proven applications in nascent RNA folding (1–4) and systematic RNA aptamer characterization (5,6). Historically, transcription arrest at a defined DNA coordinate has been achieved by attaching a protein roadblock to a DNA template to halt the progression of RNA polymerases (RNAPs). Sequence specific protein roadblocks, such the catalytically inactive EcoRI_E111Q_ mutant (7) and the *Escherichia coli* (*E. coli*) DNA replication terminator protein Tus (5), can be directed to a precise DNA location by encoding a binding sequence in the template DNA. Alternatively, the biotin-streptavidin complex is capable of halting some RNAPs if positioned either at the downstream DNA template terminus (8) or internally within the transcribed DNA strand (3). Recently, a reversible dCas9 roadblock was developed to enable time-dependent control of transcription arrest sequentially at multiple DNA positions (9). While these approaches have proven effective in diverse applications, their utility is limited to experimental contexts that tolerate the inclusion of extrinsic protein factors in the *in vitro* transcription reaction.

In contrast to protein roadblocks, chemical DNA lesions have not typically been used for nascent RNA display experiments despite the well-established inhibitory effect of DNA lesions on transcription elongation (10). The lack of chemical transcription roadblocking approaches can be attributed to two challenges: First, chemical lesions that efficiently stall RNAPs are also likely to stall DNA polymerases so that preparation of DNA templates for *in vitro* transcription yields a truncated dsDNA product during PCR amplification. Preparation of internally modified double-stranded DNA (dsDNA) has been achieved for short DNA molecules by annealing and ligating modified oligonucleotides (11) and for long DNA molecules by enzymatically generating single-strand DNA gaps that can be filled with modified oligonucleotides (12–14). However, these approaches are sequence-dependent and therefore not suitable for internally modifying the complex DNA sequence libraries that are frequently used in nascent RNA display experiments (4–6). Second, some RNAPs have been shown to bypass non-bulky DNA lesions such as abasic sites (15–19); consequently, RNAP stalling at some chemical lesions is time-dependent. In this work, we address both of the above challenges to develop a simple and versatile chemical approach for halting *E. coli* RNAP transcription elongation *in vitro*.

We have determined that an internal desthiobiotin-triethylene glycol (desthiobiotin-TEG) modification positioned in the transcribed strand of *in vitro* transcription DNA templates efficiently and stably halts *E. coli* RNAP. We first show that internally modified dsDNA can be prepared by sequential PCR amplification and translesion synthesis. We then demonstrate that desthiobiotin-TEG alone efficiently halts RNAP one nucleotide upstream of the modification site and that lesion bypass by RNAP is sufficiently slow to enable manipulation of stalled elongation complexes. Lastly, we show that transcription elongation complexes that stall at the modification site sequester desthiobiotin from streptavidin binding and are stable for at least two hours at ambient temperature. Our findings establish a method for chemically encoding a transcription stall site within a DNA molecule to enable efficient RNAP roadblocking in a minimal *in vitro* transcription reaction.

## Results

### Rationale for DNA modifier selection

We first observed that an internal desthiobiotin-TEG modification by itself (Figure 1A) in the transcribed DNA strand causes *E. coli* RNAP to stall when attempting to develop a reversible desthiobiotin-streptavidin transcription roadblock. Although the desthiobiotin-streptavidin complex was capable of halting RNAP, RNAP also appeared to stall at the desthiobiotin-TEG modification site following streptavidin elution. Given that *E. coli* RNAP has been previously reported to bypass abasic sites in the transcribed DNA strand (15), we envisioned that the desthiobiotin-TEG modification might enable protein-free transcription roadblocking at a defined DNA template position. Notably, the desthiobiotin-TEG modification is distinct from abasic lesions both due to the presence of the desthiobiotin moiety and because it introduces unnatural spacing (two vs. three carbon) in the transcribed DNA strand. We therefore performed our initial assessment of desthiobiotin-TEG as a transcription roadblock in comparison to an internal amino-linker modification, which preserves the natural three carbon spacing of the DNA phosphodiester backbone (Figure 1B).

**Figure 1.**
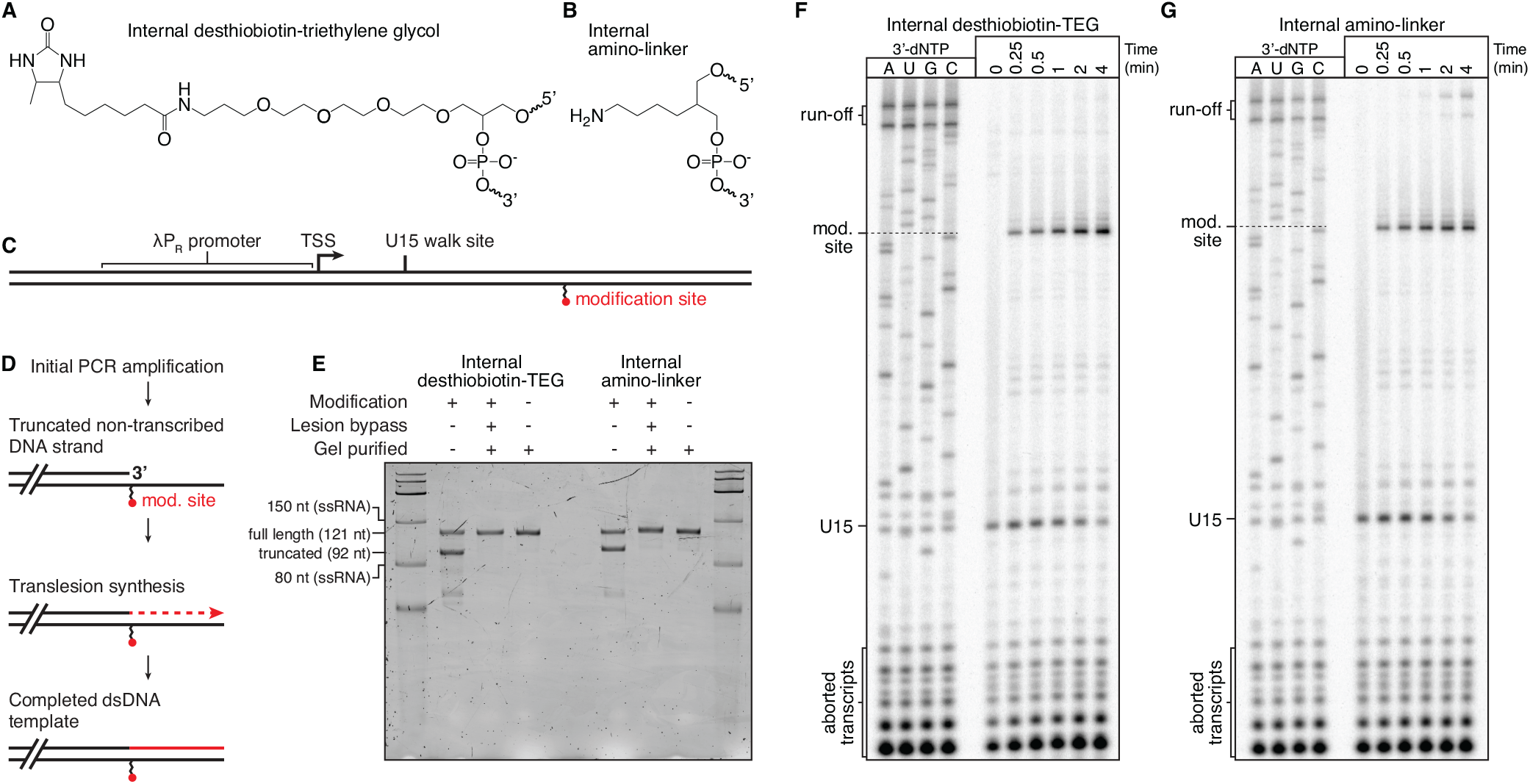
Preparation and characterization of chemical roadblocking DNA templates. **(A-B)** Chemical structures of internal desthiobiotin-TEG (A) and amino-linker (B) modifications. **(C)** Layout of internally modified DNA templates. **(D)** Overview of internally modified DNA template preparation. **(E)** Denaturing PAGE quality analysis of internally modified DNA template preparations. The size marker is the Low Range ssRNA Ladder (New England Biolabs). **(F-G)** Single-round *in vitro* transcription of internal desthiobiotin-TEG (F) and amino-linker (G) modified DNA templates. In both cases, elongation complexes stall one nucleotide upstream of the modification (mod.) site. Panel E is n=1; An equivalent quality analysis gel for the preparation of a 5’-biotinylated version of the internal desthiobiotin-TEG DNA template is shown in Supplementary Figure 1D and a second quality analysis gel of this template was performed following the completion of data collection to confirm DNA template integrity (Supplementary Figure 1C). The experiments in panels F and G were performed once to precisely map critical bands for all subsequent experiments.

### Translesion synthesis enables preparation of internally modified dsDNA

To assess the transcription roadblocking properties of the desthiobiotin-TEG and internal amino-linker modifications, we prepared *in vitro* transcription DNA templates using a synthetic oligonucleotide primer that positions the modification site 29 bp upstream of the template end (Figure 1C, Table S1). Preparation of internally modified DNA templates by PCR amplification yielded a dsDNA product with one truncated strand, suggesting that Vent Exo- DNA polymerase cannot efficiently bypass the modification site (Figure 1D, E). To complete the truncated DNA strand, we purified the initial truncated PCR product and performed primer extension using the thermostable Y-family lesion bypass polymerase *Sulfolobus islandicus* DNA polymerase IV (Dpo4) (Figure 1D). The related *Sulfolobus solfataricus* Dpo4 was shown to discriminate between correct and incorrect nucleotides with reported misincorporation frequencies between 8 × 10^−3^ and 3 × 10^−4^ (20) and Dpo4 from several *Sulfolobus* species has been used in combination with *Taq* DNA polymerase to PCR amplify damaged and ancient DNA (21). In our application, we separate PCR amplification and translesion synthesis into independent steps so that the DNA template promoter and transcribed region are synthesized first using a DNA polymerase with fidelity suitable for downstream applications. Translesion synthesis is then used to bypass the modification site and complete the truncated DNA strand. Following translesion synthesis, dsDNA templates with an internal desthiobiotin-TEG or internal amino-linker modification were indistinguishable from unmodified templates by denaturing polyacrylamide gel electrophoresis (PAGE) (Figure 1E). DNA templates containing an internal desthiobiotin-TEG modification were also analyzed by non-denaturing PAGE and showed a slight mobility shift relative to unmodified DNA (Figure S1A). DNA templates in which translesion synthesis was performed using the thermostability-enhancing dNTPs 2-amino-dATP and 5-propynyl-dCTP showed an identical mobility shift, suggesting that the shift is a consequence of the desthiobiotin-TEG modification and not instability in the DNA downstream of the modification site (Figure S1B).

### Desthiobiotin-TEG efficiently blocks *E. coli* RNAP transcription

We evaluated the transcription roadblocking properties of the internal desthiobiotin-TEG and amino-linker modifications by single-round *in vitro* transcription with *E. coli* RNAP. Transcription was initiated in the absence of CTP to walk RNAP to U15, one nucleotide upstream of the first C in the transcript, before addition of NTPs to 500 μM. Both internal modifications produced a transcription stall-site one nucleotide upstream of the modification position at C42 (Figure 1F,G). Importantly, an unmodified DNA template control showed no evidence of modification-independent transcription stalling at C42, indicating that the C42 stall site is entirely modification-dependent (Figure 2A). We next compared the desthiobiotin-TEG and amino-linker roadblocks to the ‘gold standard’ terminal biotin-streptavidin roadblock in a 32 minute time course. After 32 minutes there was no indication that RNAP had bypassed the terminal-biotin streptavidin roadblock to produce run-off transcripts (Figure 2A). In contrast to the terminal biotin-streptavidin complex, retention of elongation complexes at the desthiobiotin-TEG stall site was not absolute: after 32 minutes ~87% of elongation complexes remained at the stall site (Figure 2A, B). Nonetheless, desthiobiotin-TEG outperformed the amino-linker modification, which retained only ~40% of elongation complexes at the stall site after 32 minutes (Figure 2A, B). Lastly, we measured the rate at which elongation complexes bypass the desthiobiotin-TEG modification site following promoter escape. In our standard reaction conditions for internal RNA labeling (200 μM ATP, GTP, and CTP; 50 μM UTP), we observed an initial decay of t_1/2_=592 minutes (n=2, *R^2^* = 0.96, 95% CI [541, 655]) (Figure 3). In our first replicate, which included time points to 256 minutes, desthiobiotin-TEG bypass slowed after the 64 minute time point. In our second replicate, which included time points to 64 minutes, modification bypass slowed after the 48 minute time point (Figure 3). While the origin of this effect is unclear, it suggests that stalled elongation complexes may be heterogeneous. We conclude that virtually all elongation complexes initially stall upon encountering the desthiobiotin-TEG modification and that the vast majority of elongation complexes persist at the stall site well beyond the reaction time of typical nascent RNA display experiments.

**Figure 2.**
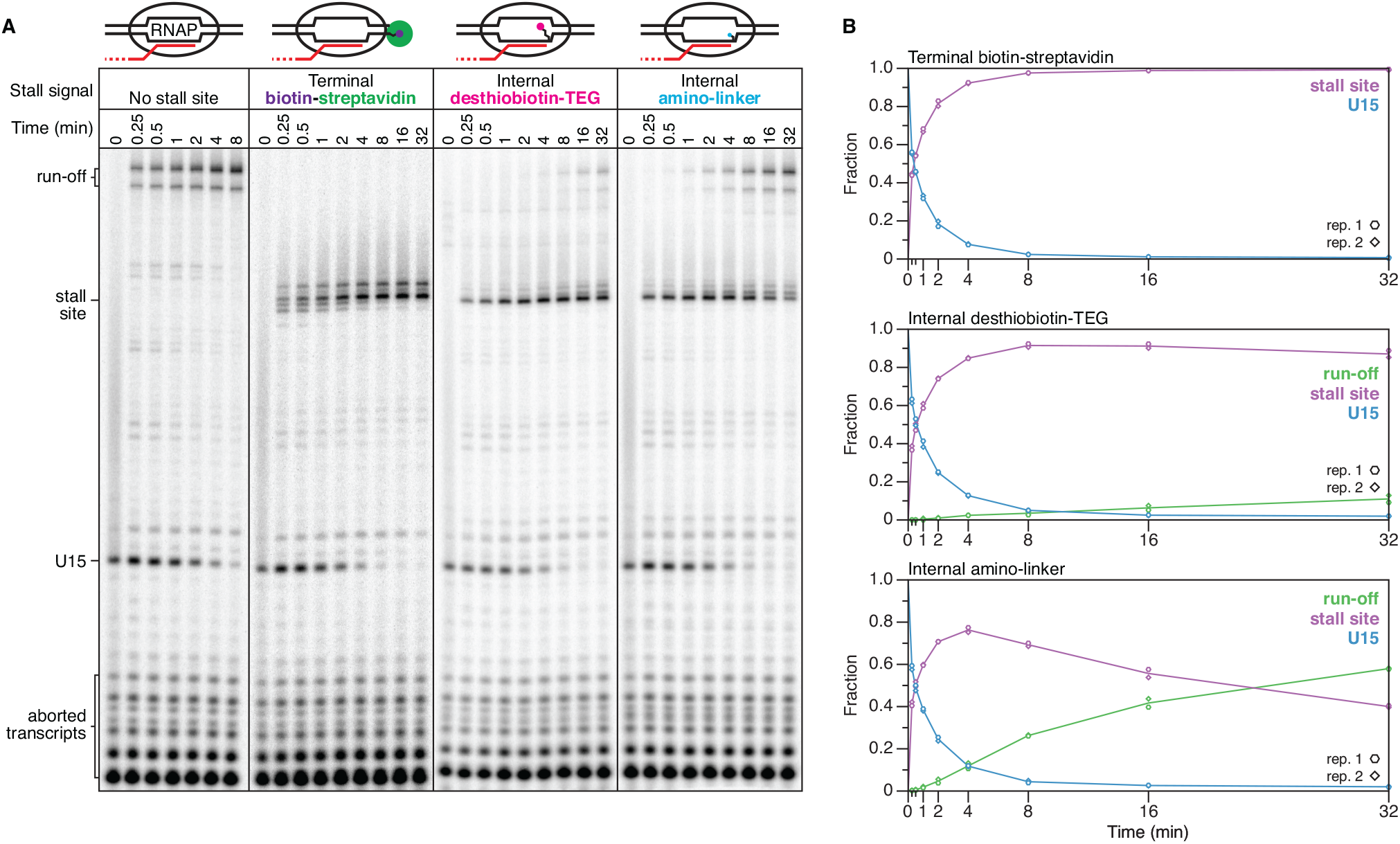
Comparison of biotin-streptavidin and chemical roadblock efficiency. **(A)** Single-round in vitro transcription of DNA templates without and with biotin-streptavidin, desthiobiotin-TEG, and amino-linker stall sites. Solid lines between gel images denote gel splices. **(B)** Quantification of gels shown in (A). All data are n=2 independent replicates. Desthiobiotin-TEG and amino-linker experiments were performed together, unmodified and biotin-streptavidin controls were performed separately.

**Figure 3.**
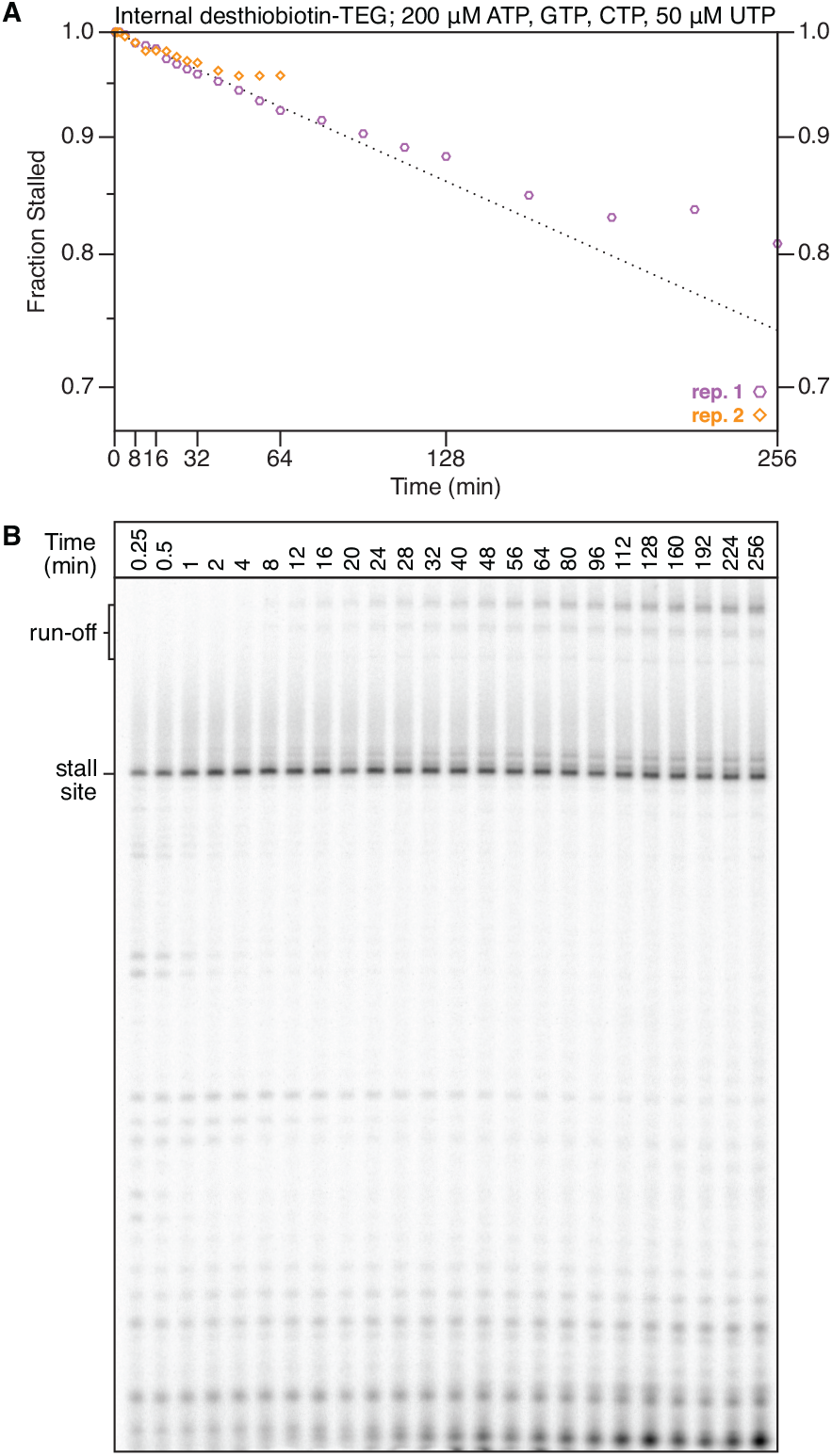
Time-dependent desthiobiotin-TEG bypass by E. coli RNA polymerase. **(A)** Plot of the fraction of elongation complexes that reached but did not bypass the desthiobiotin-TEG stall site over 256 minute (replicate 1) and 64 minute (replicate 2) time courses. The exponential decay curve was fit using time points from 2 to 48 minutes. Note that the y-axis is truncated to facilitate clear data visualization. **(B)** Gel showing the 256 minute time course from (A). Time points from 0.25 minutes to 64 minutes are n=2; time points from 80 minutes to 256 minutes are n=1.

### The *E. coli* RNAP footprint blocks streptavidin binding

The primary caveat to using the desthiobiotin-TEG modification as a transcription stall site is that it is not functionally inert. Many applications of chemical transcription roadblocking will also depend on immobilization of the DNA template, which is typically achieved by attachment of streptavidin-coated magnetic particles to an end of the template. It is therefore desirable that the DNA template contain only one attachment point. One solution to this limitation is to sequester the desthiobiotin-TEG modification within the RNAP footprint before DNA immobilization. To test this approach, we first positioned elongation complexes at the U15 walk site such that the desthiobiotin-TEG modification is exposed (Figure 4A). After incubation with streptavidin-coated magnetic particles, ~89% of U15 complexes partitioned into the bead pellet (Figure 4B). In contrast, when RNAP was positioned at the desthiobiotin-TEG stall site, only ~4% of stall site complexes were recovered in the bead fraction (Figure 4A, B). Thus, positioning *E. coli* RNAP at the desthiobiotin-TEG stall site renders the desthiobiotin moiety inert, presumably by sequestration within the RNAP active center.

**Figure 4.**
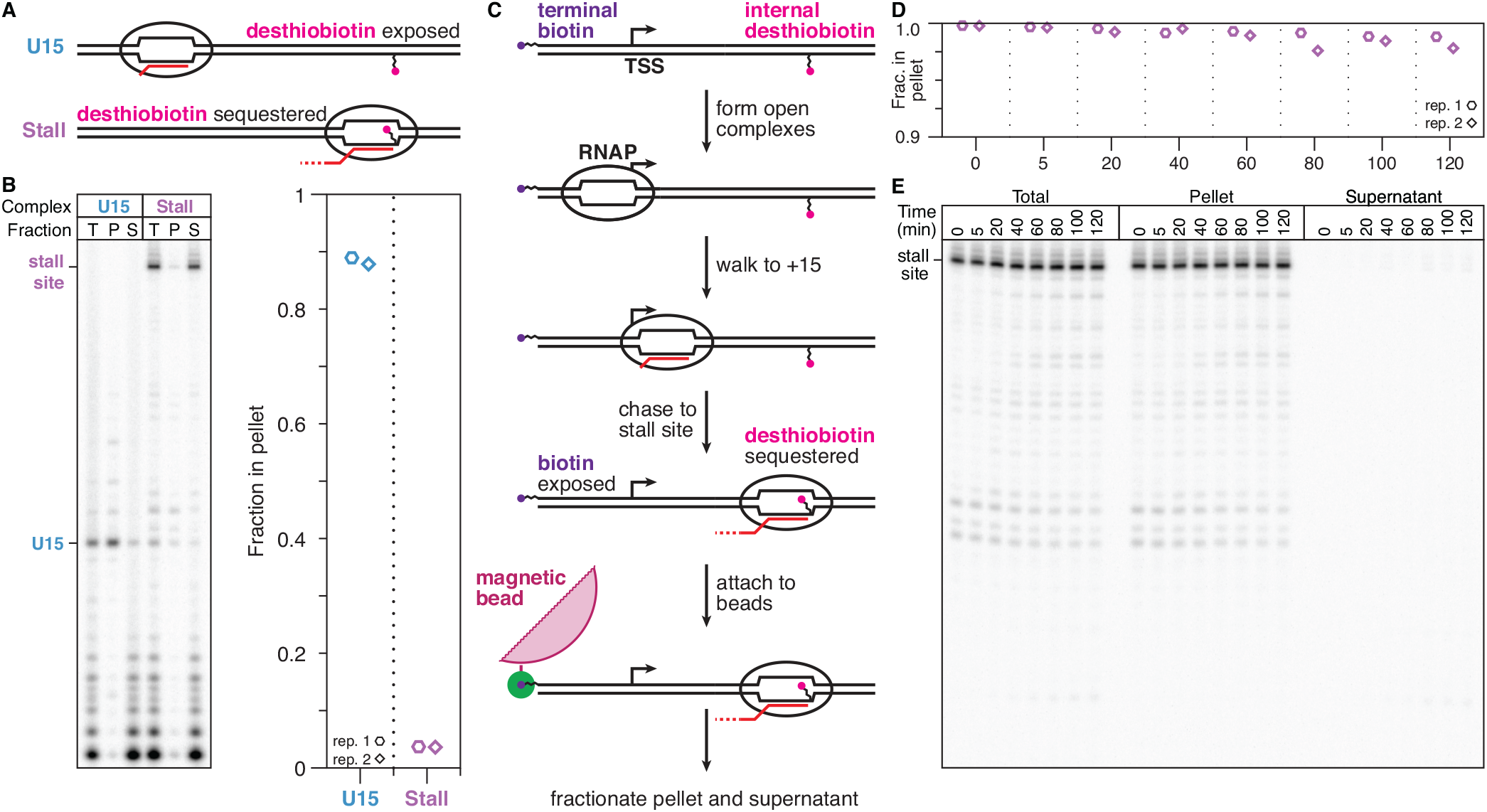
Quantification of elongation complex stability at a desthiobiotin-TEG stall site. **(A)** Overview of the desthiobiotin sequestration experiment in (B). If RNAP is positioned at U15, the desthiobiotin-TEG modification is predicted to be exposed and capable of binding streptavidin magnetic beads. If RNAP is positioned at the stall site, the desthiobiotin-TEG modification is predicted to be sequestered and incapable of binding streptavidin magnetic beads. **(B)** Fractionation of internal desthiobiotin-TEG modified DNA templates in the presence of the elongation complexes described in (A). Quantification of the pellet-bound fraction is shown. **(C)** Overview of the experiment shown in (D) and (E). Using a DNA template with an internal desthiobiotin-TEG stall site and a 5’ terminal biotin, RNAP is positioned at the stall site to sequester desthiobiotin, but 5’-biotin remains exposed. These complexes are immobilized on streptavidin magnetic beads, washed to remove free NTPs and incubated at room temperature before fractionation into bead pellet and supernatant. **(D)** Quantification of the fraction of desthiobiotin-TEG-stalled elongation complexes retained in the bead pellet over time. Note that the y-axis range is 0.9 to 1.0 to facilitate clear data visualization. **(E)** Gel showing replicate 1 from (D). Panel B is n=2. Pellet and supernatant data points in Panels D and E are n=2; the total reaction control was performed once.

### Desthiobiotin-halted elongation complexes are stably bound to DNA

The observation that the RNAP footprint sequesters desthiobiotin function enabled us to assess the stability of desthiobiotin-TEG-stalled elongation complexes over long incubation times. We prepared a DNA template containing both an internal desthiobiotin-TEG modification to stall RNAP and a terminal biotin-TEG modification upstream of the promoter for DNA immobilization by attachment to streptavidin magnetic beads (Figure 4C). After elongation complexes were positioned at the desthiobiotin-TEG stall site, the transcription reaction was incubated with streptavidin magnetic beads for 10 minutes, washed twice with transcription buffer supplemented with 1 mM MgCl_2_ to remove free NTPs, and resuspended in this same buffer (Figure 4C). ~98% of stalled elongation complexes remained attached to the magnetic particles after two hours of incubation at room temperature (Figure 4D, E). Thus, elongation complexes that are stalled at a desthiobiotin-TEG modification site retain the exceptional stability that is expected for ternary transcription elongation complexes.

## Discussion

Nascent RNA display is a powerful tool that has proven applications in mapping RNA structure and folding (1–4) and systematic characterization of RNA function (5,6). Established approaches for halting RNAP have typically relied on protein roadblocks that are not easily navigable by transcribing polymerases. While these approaches have proven effective for placing RNAP at a defined DNA template position (7–9) and for systematically distributing RNAP across every DNA template position (3), each depends on including an extrinsic protein component in the transcription reaction. Although this requirement is compatible with many applications, the chemical transcription roadblocking approach described here provides a simple alternative that expands the repertoire of nascent RNA display tools.

The primary advantage of chemical transcription roadblocking is that no extrinsic factors are needed to efficiently halt RNAP. The cost of this advantage is that, in contrast to terminal biotin-streptavidin roadblocking, transcription arrest is time-dependent: in the presence of NTPs some elongation complexes can eventually bypass the modification site (Figures 2 and 3). Nonetheless, the slow rate at which RNAP bypasses the desthiobiotin-TEG modification allows ample time for either making biochemical measurements or depleting NTPs, after which the halted ternary elongation complex persists for hours (Figure 4). Furthermore, the persistence of stalled elongation complexes for tens of minutes in the presence of 500 μM NTPs suggests that our approach is generalizable to many experimental contexts.

Our chemical approach for transcription roadblocking is enabled by the facile preparation of internally modified DNA templates of arbitrary length using translesion synthesis (Figure 1). Because this approach is sequence-independent, it is suitable both for the preparation of dsDNA templates with a defined sequence and for the preparation of complex sequence libraries. Importantly, the lesion bypass reaction is simple and efficient: in our reaction conditions translesion synthesis by *Sulfolobus* Dpo4 resulted in DNA template preparations that were indistinguishable from an unmodified template by denaturing gel electrophoresis and remained fully intact after several freeze-thaw cycles throughout data collection (Figure S1C). By performing PCR amplification and translesion synthesis sequentially, our DNA template preparation protocol enables the transcribed region of the DNA template to be amplified with fidelity that is appropriate for downstream applications. A useful feature of this two-step preparation is that removing excess dNTPs after PCR amplification enables translesion synthesis to be performed with a separate dNTP mixture (Figure S1B). Consequently, additional functionalization can be easily introduced in the non-transcribed DNA strand downstream of the chemical roadblock by inclusion of modified dNTPs during translesion synthesis.

While our study clearly establishes the desthiobiotin-TEG modification as a potent transcription blockade, it remains unclear why this modification stalls RNAP more efficiently than abasic lesions (15) or an amino-linker modification (Figure 2). One possible mechanism is that the branched triethylene glycol linker that connects the desthiobiotin moiety to the template DNA interrupts the phosphodiester backbone with a two carbon insertion as opposed to the natural three carbon spacing of the deoxyribose sugar. A second possibility is that the desthiobiotin-TEG structure itself interferes with catalysis of nucleotide addition by RNAP. It is likely that other readily available oligonucleotide modifications are suitable for chemical transcription roadblocking. For example, etheno-dA was recently shown to halt *E. coli* RNAP efficiently in the presence of 100 μM NTPs (19). We suggest that a DNA modification is suitable for chemical transcription roadblocking if it satisfies four criteria: 1) the modification should efficiently halt the target RNAP; 2) the modification should be efficiently bypassed by a translesion DNA polymerase to facilitate DNA template preparation; 3) the modification should be chemically stable; and 4) the modification should not interfere with downstream applications. We envision that the chemical transcription roadblocking approach presented in this work will facilitate increasingly demanding nascent RNA display applications by enabling RNA synthesis in a minimal transcription reaction.

## Experimental procedures

### Oligonucleotides

All oligonucleotides were purchased from Integrated DNA Technologies (Coralville, IA) and are described in Table S1. All modified oligonucleotides were HPLC-purified to ensure efficient modification. The oligonucleotide used as a template for PCR amplification was PAGE-purified to enrich for complete product.

### Unmodified DNA template preparation

Linear DNA templates without internal modifications were prepared by PCR amplification essentially as described previously (3). Briefly, five 100 μl reactions containing 81.5 μl of water, 10 μl Thermo Pol Buffer (New England Biolabs, Ipswich, MA), 2 μl of 10 mM dNTPs (New England Biolabs), 2.5 μl of 10 μM oligonucleotide A (unmodified forward primer; Table S1), 2.5 μl of 10 μM oligonucleotide C (unmodified reverse primer; Table S1) or oligonucleotide F (5’ biotinylated reverse primer, Table S1), 1 μl of Vent Exo- DNA polymerase (New England Biolabs), and 0.5 μl of 0.1 nM oligonucleotide G (template oligonucleotide, Table S1) were amplified for 30 PCR cycles. Following amplification, 100 μl reactions were combined into two 250 μl pools and precipitated by adding 25 μl of 3M sodium acetate (NaOAc) pH 5.5 and 750 μl of cold 100% ethanol (EtOH), chilling at −80C for 15 minutes, and centrifugation at 20,000 x g for 15 minutes. DNA pellets were washed with 70% EtOH (v/v), dried using a SpeedVac, dissolved in 30 μl of nuclease-free water, run on a 1% (wt/v) agarose gel, and extracted using a QIAquick Gel Extraction Kit (Qiagen, Hilden, Germany) according to the manufacturer’s protocol. DNA concentration was determined by a Qubit 1.0 Fluorometer (Life Technologies, Carlsbad, CA). The fully assembled DNA template sequence is shown in Table S1.

### Internally modified DNA template preparation

Internally modified linear DNA templates were PCR amplified as above except that oligonucleotides A or B (unmodified and 5’ biotinylated forward primer, respectively, Table S1) were used as a forward primer, oligonucleotides D or E (internal desthiobiotin-TEG and internal amino-linker reverse primers, respectively, Table S1) were used as a reverse primer, and eleven 100 μl reactions were prepared. Following PCR amplification, each 100 μl reaction was purified using a QIAquick PCR Purification Kit (Qiagen) according to the manufacturer’s protocol and eluted in 30 μl of nuclease-free water. Of the eleven 30 μl aliquots, one was saved as a control for DNA template quality analysis and ten were pooled for translesion synthesis, quantified using a Qubit 1.0 Fluorometer (Life Technologies), and total volume was determined; typical yield at this stage was approximately 4.5 μg of DNA. Translesion synthesis reactions were prepared by combining the eluted DNA with 100 μl of Thermo Pol Buffer (New England Biolabs), 20 μl of 10 mM dNTPs (New England Biolabs), 10 μl of *Sulfolobus* DNA polymerase IV (New England Biolabs) and nuclease-free water to 1 ml. The 1 ml master mix was split into 100 μl aliquots and incubated at 55C for 1 hour. Translesion synthesis with thermostability enhancing dNTPs was performed as above, but with a dNTP mixture in which ATP and CTP were completely substituted with 2-amino-dATP and 5-propynyl-dCTP (TriLink Biotechnologies, San Diego, CA). Template DNA was then precipitated, gel extracted, and quantified as described for unmodified DNA templates.

### DNA template quality control

To verify success of the translesion synthesis reaction, all internally modified DNA templates were subjected to quality control by denaturing Urea polyacrylamide gel electrophoresis (PAGE) using the UreaGel System (National Diagnostics, Atlanta, GA) (Figure 1E, Figure S1C, D). Each quality control gel included a reaction aliquot that had not undergone translesion synthesis as a negative control and an unmodified positive control. A 10% polyacrylamide urea-PAGE gel was pre-run at 300 V (Figure 1E and Figure S1D) or 250 V (Figure S1C) for 30 minutes on a Mini-PROTEAN Tetra Cell that was assembled so that 1x Tris/Borate/EDTA (TBE) buffer in the outer chamber covered only the bottom ~1 centimeter of the gel plates. Immediately prior to loading, 1 μl of 50 nM DNA template (50 fmol) was mixed with 15 μl of formamide loading dye (1x transcription buffer (described below under *Single-round in vitro transcription*), 80% (v/v) formamide, 0.025% (wt/v) bromophenol blue and xylene cyanol) and boiled for 5 minutes before placing on ice for 2 minutes. All non-denaturing PAGE quality control gels were 8% polyacrylamide. Gels were stained with SYBR Gold (Life Technologies) and imaged using an Amersham Biosciences Typhoon 9400 Variable Mode Imager. Following collection of all data, a second denaturing PAGE quality control gel was performed to confirm that the internal desthiobiotin modified DNA template remained fully intact (Figure S1C).

### Single-round *in vitro* transcription

Single-round *in vitro* transcription reactions were performed essentially as previously described (4,22). All single-round transcription reactions contained 5 nM DNA template and 0.016 U/μl *E. coli* RNA polymerase holoenzyme (New England Biolabs) in transcription buffer (20 mM tris(hydroxymethyl)aminomethane hydrochloride (Tris-HCl) pH 8.0, 0.1 mM ethylenediaminetetraacetic acid (EDTA), 1 mM dithiothreitol (DTT) and 50 mM potassium chloride (KCl)), and 0.2 mg/ml bovine serum albumin (BSA). All NTP mixtures were prepared using High Purity ATP, GTP, CTP, and UTP (GE Life Sciences, Chicago, IL). Two protocol variations were performed: For experiments in which transcription was initiated by walking elongation complexes to +15 relative to the transcription start site, elongation complexes were stalled at U15 by incubating reactions containing 2.5 μM ATP and GTP, 1.5 μM UTP, 0.2 μCi/μl [α-^32^P]UTP (Perkin-Elmer, Waltham, MA), and 10 mM MgCl_2_ at 37C for 10 minutes before adding NTPs to 500 μM and rifampicin (Gold Biotechnology, St. Louis, MO) to 10 μg/ml. RNA sequencing ladders were generated by walking elongation complexes to +15 before adding NTPs supplemented with a chain terminating 3’-deoxyNTP (TriLink Biotechnologies) to 100 μM; reactions proceeded at 37C for 5 minutes before addition of 125 μL of stop solution. For Figure 2, the biotin-streptavidin roadblock was prepared by adding streptavidin monomer (Promega, Madison, WI) to 100 nM. For Figure 3, open promoter complexes were formed by incubating reactions containing 200 μM ATP, GTP, CTP, 50 μM UTP and 0.2 μCi/μl [α-^32^P]UTP (Perkin-Elmer) at 37C for 10 minutes and initiated by adding magnesium chloride (MgCl_2_) to 10 mM and rifampicin to 10 μg/ml. A mineral oil overlay was applied to the reaction master mix after taking the 4 minute time point to prevent evaporation over the 256 minute and 64 minute time courses. Time courses were performed at 37C; at each time point, one 25 μl reaction volume was added to 125 μL of stop solution (0.6 M Tris, pH 8.0, 12 mM EDTA). All reactions were extracted by adding 150 μL of phenol/chloroform/isoamyl alcohol (25:24:1), vortexing, centrifugation, and collection of the aqueous phase and then ethanol precipitated by adding 450 μL of 100% ethanol, 1.2 μl of GlycoBlue Coprecipitant (Thermo Fisher Scientific, Waltham, MA) and storing at −20C overnight. After centrifugation and removal of bulk and residual ethanol, precipitated RNA was resuspended in formamide loading dye and fractionated by Urea-PAGE using a 12% polyacrylamide sequencing gel prepared with the UreaGel System (National Diagnostics, Atlanta, GA).

### Equilibration of streptavidin magnetic beads

All transcription reactions with magnetic separation were performed using 10 μl of Dynabeads MyOne Streptavidin C1 beads (Invitrogen, Waltham, MA) per 25 μl transcription volume. For each experiment, magnetic beads were prepared in bulk by removing storage buffer, incubating with 500 μl hydrolysis buffer (100 mM sodium hydroxide, 50 mM sodium chloride (NaCl)) for 10 minutes at room temperature with rotation, washing once with 1 ml of high salt wash buffer (50 mM Tris-HCl pH 7.5, 2 M NaCl, 0.5% Triton X-100), once with 1 ml of binding buffer (10 mM Tris-HCl pH 7.5, 300 mM NaCl, 0.1% Triton X-100), twice with 1 ml of 1X transcription buffer supplemented with Triton X-100 to 0.1%, and resuspended in 1x transcription buffer with 0.1% Triton X-100 to a volume equivalent to that of the total transcription master mix for storage until use.

### Desthiobiotin protection assay

Transcription was initiated by walking elongation complexes to U15 as described above. To assess desthiobiotin-streptavidin binding when RNAP is at U15, one half of the reaction was mixed with rifampicin to 10 μg/mL and immediately mixed with streptavidin magnetic beads. To assess desthiobiotin-streptavidin binding when RNAP is at the desthiobiotin-TEG stall site, the second half of the reaction was mixed with NTPs to 500 μM and rifampicin to 10 μg/ml and incubated at 37C for 5 minutes before mixing with streptavidin magnetic beads. After 10 minutes of incubation, 25 μl of the reaction was added to 125 μl stop solution as a ‘total’ sample control and an additional 25 μl was added to a new tube and placed on a magnetic stand for 1 minute to separate the bead pellet and supernatant. The supernatant was added to 125 μl of stop solution and the pellet was resuspended in 25 μl of 1x transcription buffer supplemented with 10 mM MgCl_2_ before being added to 125 μl of stop solution. Samples were phenol/chloroform extracted, precipitated, and fractionated by Urea-PAGE as described above.

### Elongation complex stability assay

Elongation complex stability assays were performed using a DNA template containing both an internal desthiobiotin-TEG modification and a terminal biotin-TEG modification. Stalled elongation complexes were prepared by first walking elongation complexes to U15 as described above and then chasing RNAP to the desthiobiotin-TEG stall site by adding NTPs to 500 μM and rifampicin to 10 μg/mL and incubating at 37C for 5 minutes. The transcription reaction was then mixed with pre-equilibrated streptavidin magnetic beads and incubated at 25C for 10 minutes. Following DNA template immobilization, beads were washed twice with 1 ml of 1x transcription buffer supplemented with 1 mM MgCl_2_ to remove free NTPs. After removing a ‘zero’ time point for fractionation, the transcription reaction was incubated at room temperature for 2 hours with rotation and time points were taken as indicated. At each time point, 25 μl of the reaction was added to 125 μl stop solution as a ‘total’ sample control and a second 25 μl was added to a new tube and placed on a magnetic stand for 1 minute to separate the bead pellet and supernatant. The supernatant was added to 125 μl of stop solution and the pellet was resuspended in 25 μl of 1x transcription buffer supplemented with 1 mM MgCl_2_ before being added to 125 μl of stop solution. Samples were phenol/chloroform extracted, precipitated, and fractionated by Urea-PAGE as described above.

## Quantification of radiolabeled *in vitro* transcription reactions

Reactive nucleotides were detected by an Amersham Biosciences Typhoon 9400 Variable Mode Imager and quantified using ImageQuant (GE Life Sciences). For experiments in which elongation complexes were walked to U15, transcripts were considered end-labeled due to the high probability of radiolabel incorporation during the initial walk (~4.25% per U nucleotide) and the low probability of radiolabel incorporation during the chase (~0.013% per U nucleotide) and no normalization was applied. For the experiments in Figure 3, band intensity was divided by transcript U content to normalize for the incorporation of [α-^32^P]UTP. In Figure 2, the fraction of run-off, stall site, and U15 transcriptions was determined by dividing the band intensity of each transcript class by the total band intensity of all three classes. In Figure 3, fraction stalled was determined by dividing the normalized band intensity of desthiobiotin-TEG-stalled transcripts by sum of the normalized band-intensity of stalled and run-off transcripts; t_1/2_ was determined by applying a one phase exponential decay fit to time points from 2 minutes (after new arrival of elongation complexes at the stall site is negligible) to 48 minutes (before the semi-logarithmic plot deviates from a straight line) (23) using GraphPad Prism 8. In Figure 4, the fraction of transcripts in the bead pellet was determined by dividing the band intensity of the indicated complex in the pellet fraction by the sum of the band intensities for the indicated complex in both the pellet and supernatant fractions.

## Acknowledgements

We thank Jeffrey W. Roberts for thoughtful discussions and critical reading of the manuscript. This work was supported by an Arnold O. Beckman Postdoctoral Fellowship (to E.J.S.), by Grant Number GM25232 from the National Institute of General Medical Sciences of the National Institutes of Health (to J.T.L)., and by Searle Funds at The Chicago Community Trust (to J.B.L.). The content is solely the responsibility of the authors and does not necessarily represent the official views of the National Institutes of Health.

## Conflict of interest

The authors declare that they have no conflicts of interest with the contents of this article.

## Author contributions

Conceptualization, E.J.S.; Methodology, E.J.S.; Investigation, E.J.S.; Formal Analysis, E.J.S.; Validation, E.J.S.; Writing – Original Draft, E.J.S.; Writing – Review & Editing, E.J.S., J.T.L, and J.B.L.; Funding Acquisition, E.J.S., J.T.L, and J.B.L.

## Materials Included

**Figure S1.**
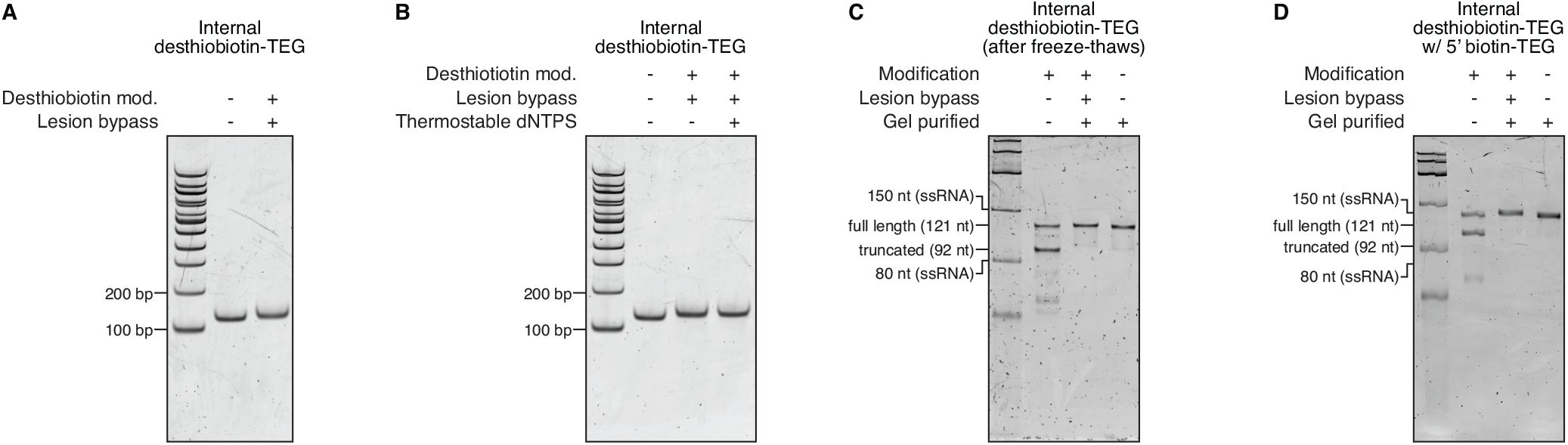
Additional DNA template quality analyses. **(A)** Non-denaturing PAGE quality analysis of DNA template with an internal desthiobiotin-TEG modification. The size marker is the Quick-Load 100 bp DNA Ladder (New England Biolabs). **(B)** Non-denaturing PAGE of DNA templates in which the initial PCR amplification was split to perform translesion synthesis with either standard dNTPs or a thermostable dNTP mixture in which dATP and dCTP were substituted with 2-amino-dATP and 5-propynyl-dCTP. The size marker is the Quick-Load 100 bp DNA Ladder (New England Biolabs). **(C)** Denaturing PAGE quality analysis of DNA template with an internal desthiobiotin-TEG modification that was performed after several freeze-thaw cycles that occurred over the course of data collection. The size marker is the Low Range ssRNA Ladder (New England Biolabs). **(D)** Denaturing PAGE quality analysis of DNA template with an internal desthiobiotin-TEG modification and a 5’ biotin-TEG modification. The size marker is the Low Range ssRNA Ladder (New England Biolabs).

**Table S1.**
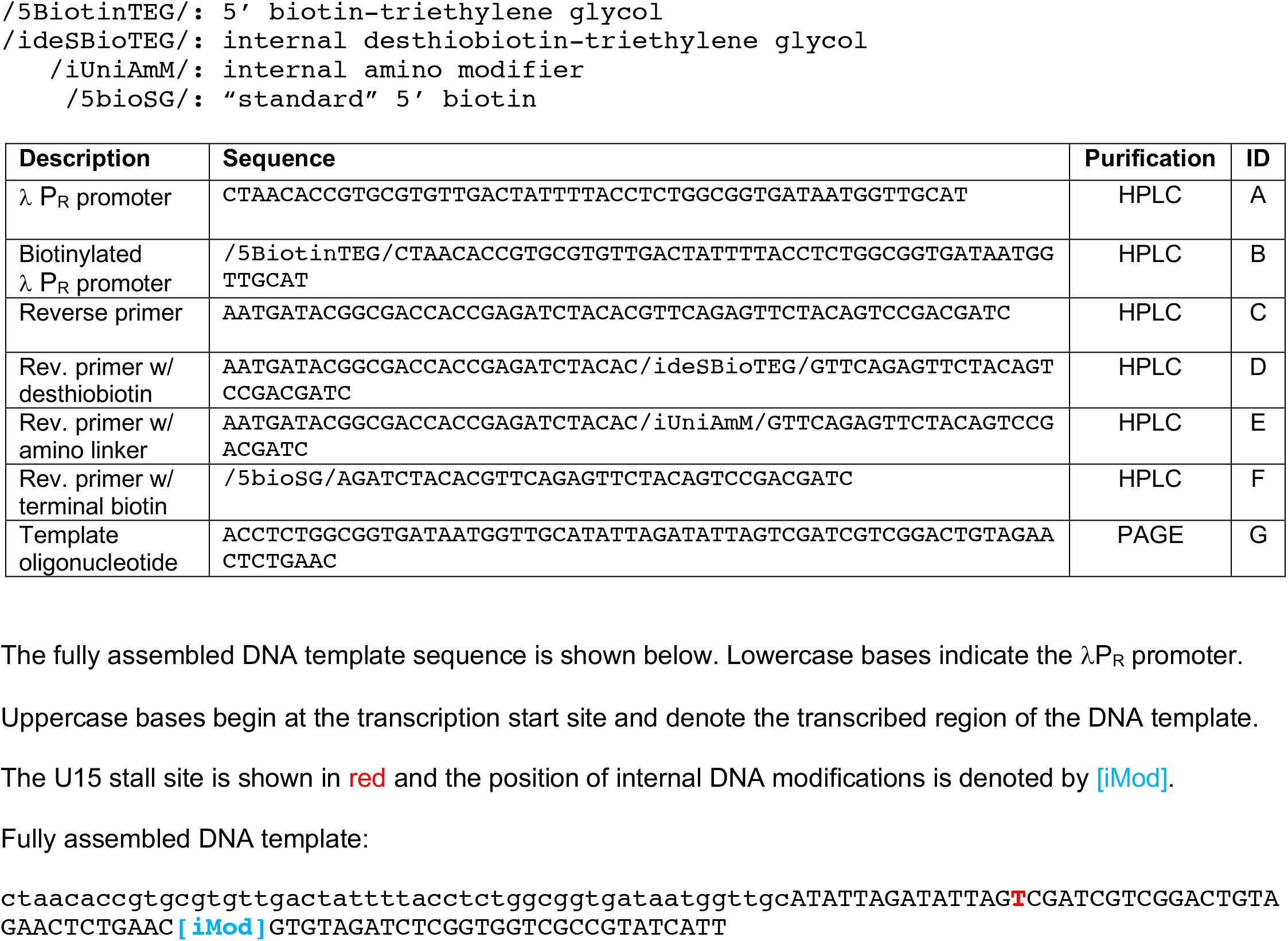
Oligonucleotides used for DNA template amplification. Below is a table of oligonucleotides used for the preparation of *in vitro* transcription DNA templates. The modification codes defined below are used for compatibility with the Integrated DNA Technologies ordering.

## Notes

#### Summary of Updates

1) Added results section 'Rationale for DNA modifier selection.' 2) Replaced gel images in Figure 1E and Figure S1D with technical replicates in which denaturing PAGE was performed at a higher voltage to improve dsDNA denaturation. 3) Added DNA sequence information for completely assembled dsDNA templates in Table S1.

